# KnetMiner: a comprehensive approach for supporting evidence-based gene discovery and complex trait analysis across species

**DOI:** 10.1101/2020.04.02.017004

**Authors:** Keywan Hassani-Pak, Ajit Singh, Marco Brandizi, Joseph Hearnshaw, Sandeep Amberkar, Andrew L. Phillips, John H. Doonan, Chris Rawlings

## Abstract

Generating new ideas and scientific hypotheses is often the result of extensive literature and database reviews, overlaid with scientists’ own novel data and a creative process of making connections that were not made before. We have developed a comprehensive approach to guide this technically challenging data integration task and to make knowledge discovery and hypotheses generation easier for plant and crop researchers. KnetMiner can digest large volumes of scientific literature and biological research to find and visualise links between the genetic and biological properties of complex traits and diseases. Here we report the main design principles behind KnetMiner and provide use cases for mining public datasets to identify unknown links between traits such grain colour and pre-harvest sprouting in *Triticum aestivum*, as well as, an evidence-based approach to identify candidate genes under an *Arabidopsis thaliana* petal size QTL. We have developed KnetMiner knowledge graphs and applications for a range of species including plants, crops and pathogens. KnetMiner is the first open-source gene discovery platform that can leverage genome-scale knowledge graphs, generate evidence-based biological networks and be deployed for any species with a sequenced genome. KnetMiner is available at http://knetminer.org.

## INTRODUCTION

Genomics is undergoing a revolution. Unprecedented amounts of data are being generated to gain deeper insight into the complex nature of many traits and diseases (Boyle et al., 2017; Stephens et al., 2015). The growing landscape of diverse and interconnected data can often hinder scientists from translating complex and sometimes contradictory information into biological understanding and discoveries. Searching for information can quickly become complex and time-consuming, which is prone to information being overlooked and subjective biases being introduced. Even when the task of gathering information is complete, it is demanding to assemble a coherent view of how each piece of evidence might come together to “tell a story” about the biology that can explain how multiple genes might be implicated in a complex trait or disease. New tools are needed to provide scientists with a more fine-grained and connected view of the scientific literature and databases, rather than the conventional information retrieval tools currently at their disposal.

Scientists are not alone with these challenges. Search systems form a core part of the duties of many professions. Studies have highlighted the need for search systems that give confidence to the professional searcher and therefore trust, explainability, and accountability remain a significant challenge when developing such systems (Russell-Rose et al., 2018). The amount of time spent on a task also influences human choice about whether to continue the task (Sweis et al., 2018). When implemented well, search systems can give a head start to researchers by cutting the time and cost to review genes, traits or molecules of interest before initiating expensive experiments. Additionally, they offer a framework for the prioritization of future research, which can highlight gaps in knowledge.

Knowledge graphs (KG) are increasingly used to make search and information discovery more efficient (Fensel et al., 2020). KGs are contributing to various Artificial Intelligence (AI) applications including link prediction, node classification, and recommendation and question answering systems (Ali et al., n.d.; Sheth et al., 2019). KGs model heterogeneous knowledge domains by integrating information into advanced unified data schemas (i.e. ontologies) and leverage that to apply formal and statistical inference methods to derive new knowledge (Ehrlinger & Wöß, 2016). Compared to more traditional data models, knowledge graphs are very flexible at integrating and searching connected heterogeneous data, where data schemas are not established a-priori (Yoon et al., 2017), and often subject to frequent changes. KGs in various forms have been widely adopted in many disciplines, ranging from social sciences to engineering, physics, computer science, design and manufacturing. Different research labs, including ourselves, are building biological KGs aimed at supporting crop improvement (Hassani-Pak et al., 2016; Xiaoxue et al., 2019), drug-target discovery (Mohamed et al., 2019), and disease-gene prioritization (Alshahrani & Hoehndorf, 2018; Messina et al., 2018).

The integrated, semi-structured and machine readable nature of KGs provides an ideal basis for the development of sophisticated knowledge discovery and data mining (KDD) applications (Holmes, 2014; Sacchi & Holmes, 2016). Exploratory data mining (EDM), a sub discipline of knowledge discovery, requires an extensive exploration stage, using both intelligent and intuitive techniques, before predictive modelling and confirmatory analysis can realistically and usefully be applied (De Bie, 2013; De Bie & Spyropoulou, 2013). Furthermore, it is considered important to include the end user into the “interactive” knowledge discovery process with the goal of supporting human intelligence with artificial intelligence (Holzinger & Jurisica, 2014). Several reports have described the benefits attained by leveraging the unique human cognitive capabilities we have, both within the fields of pattern recognition and higher-order reasoning, to interpret complex biological data and help extract biologically meaningful interpretations (Isenberg et al., 2013; Lee et al., 2012). Visualising biological information in a concise format and user-centred design can help achieve this (Fox & Hendler, 2011; Pavelin et al., 2012).

There are, however, a few important research challenges that need resolving before KDD and EDM techniques can optimally be applied to KGs. These include the formalisation of concepts such as an ‘interesting pattern’ found in a genome-scale KG, since ‘interestingness’ is subjective and will depend on the user’s perspective. The concept of ‘explaining a specific biological story’ using a minimum set of non-redundant and relevant patterns from the KG also needs to be formalised. These theoretical insights need to be turned into useful, scalable and interactive tools, suitable for use by non-experts and tested against real biological problems.

We have previously described our approaches to build genome-scale KGs (Hassani-Pak et al., 2016), to extend KGs with novel gene-phenotype relations from the literature (Hassani-Pak et al., 2010), to publish KGs as standardised and interoperable data based on FAIR principles (Brandizi et al., 2018a) and to visualise biological knowledge networks in an interactive web application (Singh et al., 2018). Our data integration approach to build KGs is based on an intelligent data model with just enough semantics to capture complex biological relationships between genes, traits, diseases and many more information types derived from curated or predicted information sources (Figure 1). In this paper, we describe the KnetMiner knowledge discovery platform (knetminer.org) for searching large genome-scale KGs and visualising interesting subgraphs of connected information about the biology of traits and diseases. KnetMiner is customizable and portable and therefore provides a cost-effective delivery platform for application to new species. We provide use-cases to demonstrate how KnetMiner has helped scientists to tell the story of complex traits and diseases in *Arabidopsis thaliana* and *Triticum aestivum* (bread wheat). The methods section describes the algorithms behind core discovery features of KnetMiner, i.e. identifying interesting subgraphs and using these to rank candidate genes.

**Figure 1:**
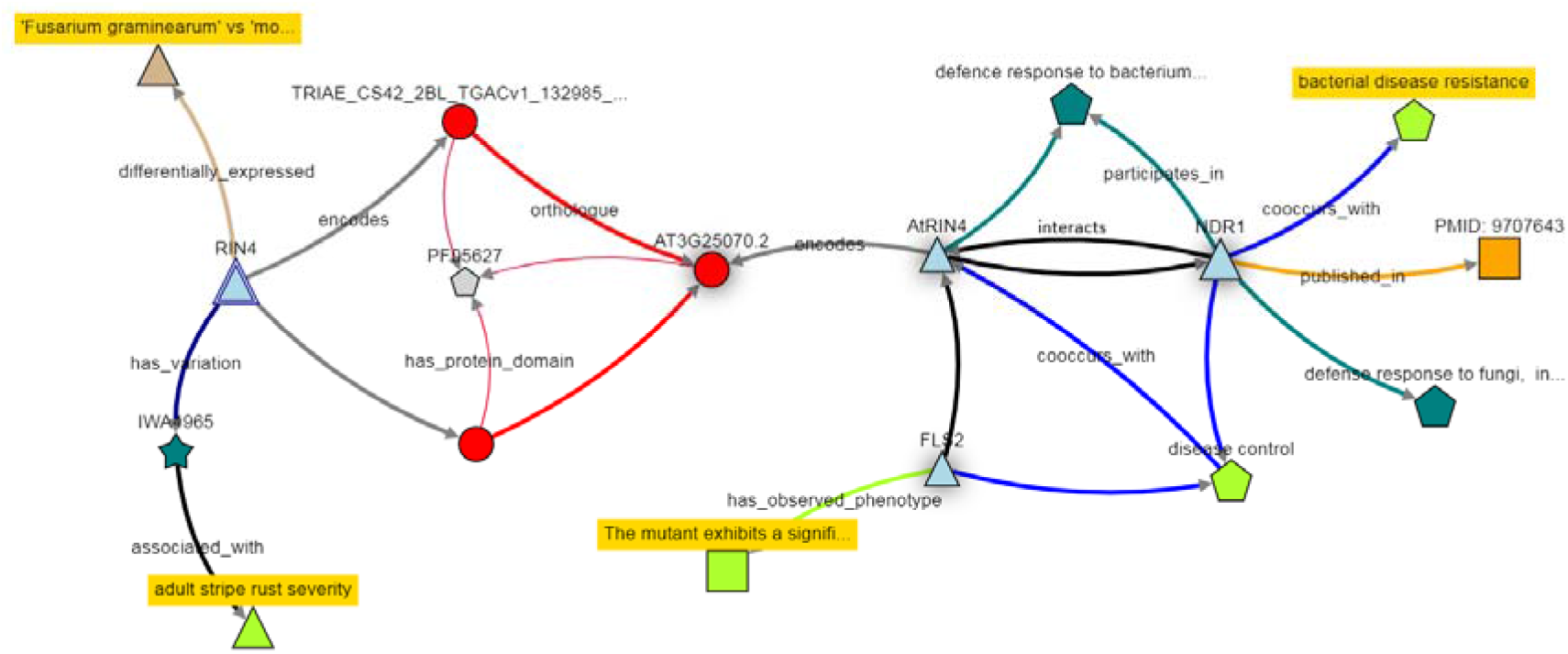
Extract of information available in the KnetMiner Knowledge Graph.

## CASE STUDIES

KnetMiner can assist in various stages of a typical research and discovery project: from early stages of literature review and hypothesis generation to later stages of biological understanding and hypothesis validation. The user-centric web interfaces have been designed to provide effective user journeys for the exploration of complex connected data. A simple search interface triggers a sophisticated search process and takes the user in two steps through a rich knowledge discovery experience (Figure 2). We have selected two biological case studies that show the application of KnetMiner in gene-trait discovery and candidate gene prioritization in a model and non-model species.

**Figure 2:**
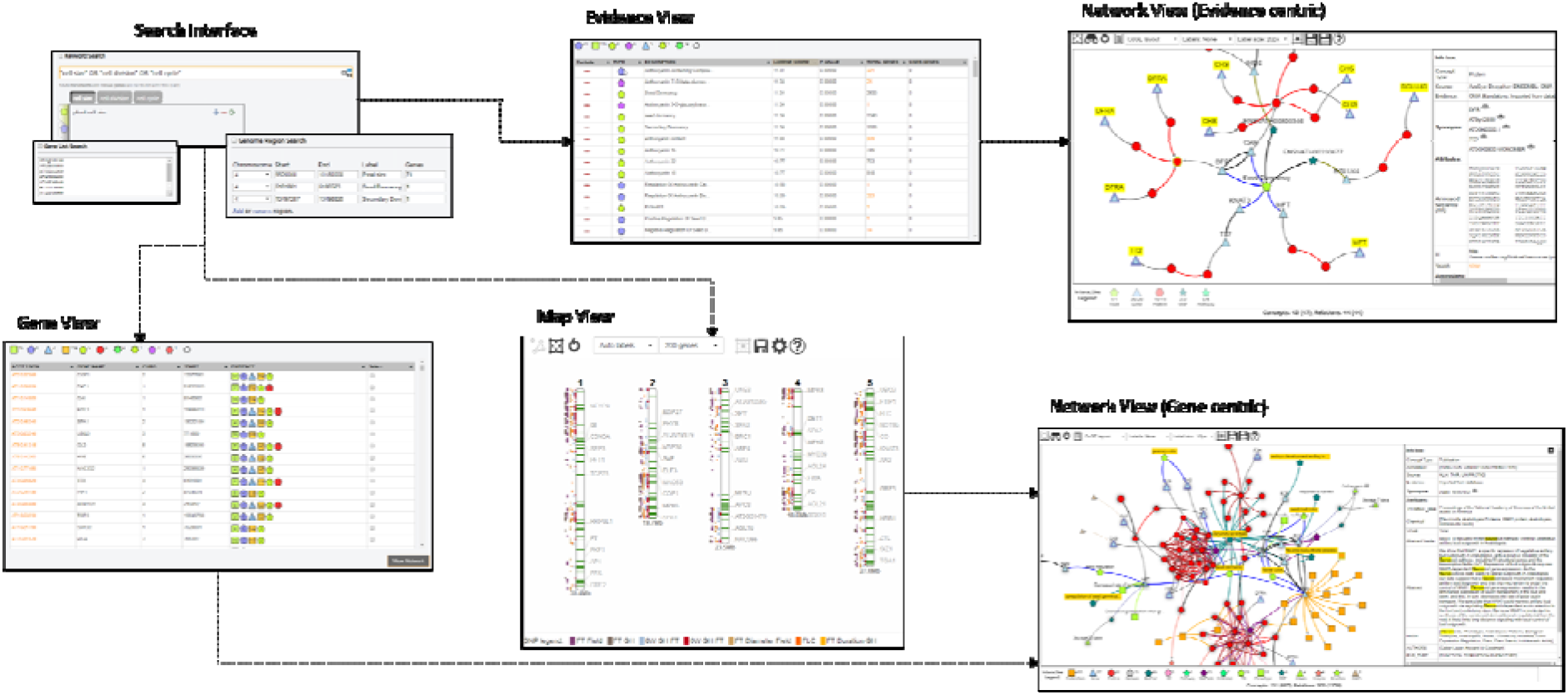
User journeys in KnetMiner. Users start with a search for keywords, genes and regions. KnetMiner provides search term suggestions and real-time query feedback. From a search, a user is presented with the following views: ***Gene View*** is a ranked list of candidate genes along with a summary of related evidence types. ***Map View*** is a chromosome based display of QTL, GWAS peaks and genes related to the search terms. ***Evidence View*** is a ranked list of query related evidence terms and enrichment scores along with linked genes. By selecting one or multiple elements in these three views, the user can get to the ***Network View*** to explore a gene-centric or evidence-centric knowledge network related to their query and the subsequent selection.

### Gene-trait discovery

KnetMiner is being used extensively to drive gene-trait discovery research in the publicly funded Designing Future Wheat programme (https://designingfuturewheat.org.uk/), see for example (Adamski et al., 2020; Alabdullah et al., 2019; Harrington et al., 2019). Wheat (*Triticum aestivum*) is the third most-grown cereal crop in the world after maize and rice, and has a hexaploid 15 Gb genome which is 5 times the size of the human genome (The International Wheat Genome Sequencing Consortium (IWGSC) et al., 2018). White-grained wheat varieties lack the red compounds (flavonoids) of the seed coat and are milder in flavor. However, white grains are prone to pre-harvest sprouting (PHS) which causes the grain to germinate before harvest and results in a loss of grain quality. It has been known for some time that PHS is associated with grain colour (Nilsson-Ehle, 1914) and that the red pigmentation of wheat grain is controlled by *R* genes on the long arms of chromosomes 3A, 3B, and 3D (Sears, 1944). However, after decades of research, it still remains unclear whether there is a potential link between the grain color gene R (Myb) and other phenotypes such as PHS.

We used KnetMiner to search for <u>TRAESCS3D02G468400</u> - the wheat *R* gene (the orthologue of Arabidopsis *TT2*) on chromosome 3D, and to explore its knowledge network generated by KnetMiner. The *TT2* network has a total of 823 connected nodes of 11 different types (see Supp Table 1) including wheat specific information sources but also cross-species information from model organisms such as Arabidopsis and rice. Similarly a range of relation types are present in the network including homologies, transcription factor target relations, protein protein interactions, phenotypic observations and correlations from mutant and genetic studies, as well as, curated or auto generated links to ontology terms and publications.

Prior to visualising the network, KnetMiner applies a graph filter for interesting subgraphs which uses the keywords that were provided as part of the search (see Methods - Graph Interestingness). In our case, since the *TT2* gene search was performed without additional keywords, a default filter is applied which hides all paths but those containing traits and phenotypes. This reduces the network from 823 nodes down to 245 nodes including 101 Trait, 48 Phenotype, 72 SNP, 22 Gene and 2 Protein nodes (Figure 3A). This network is displayed in the Network View which provides interactive features to hide or add specific evidence types from the network. Nodes are displayed in a defined set of shapes, colors and sizes to distinguish different types of evidence. A shadow effect on nodes indicates that more information is available but has been hidden. The auto-generated network, however, is not yet telling a story that is specific to our traits of interest and is limited to evidence that is phenotypic in nature.

**Figure 3:**
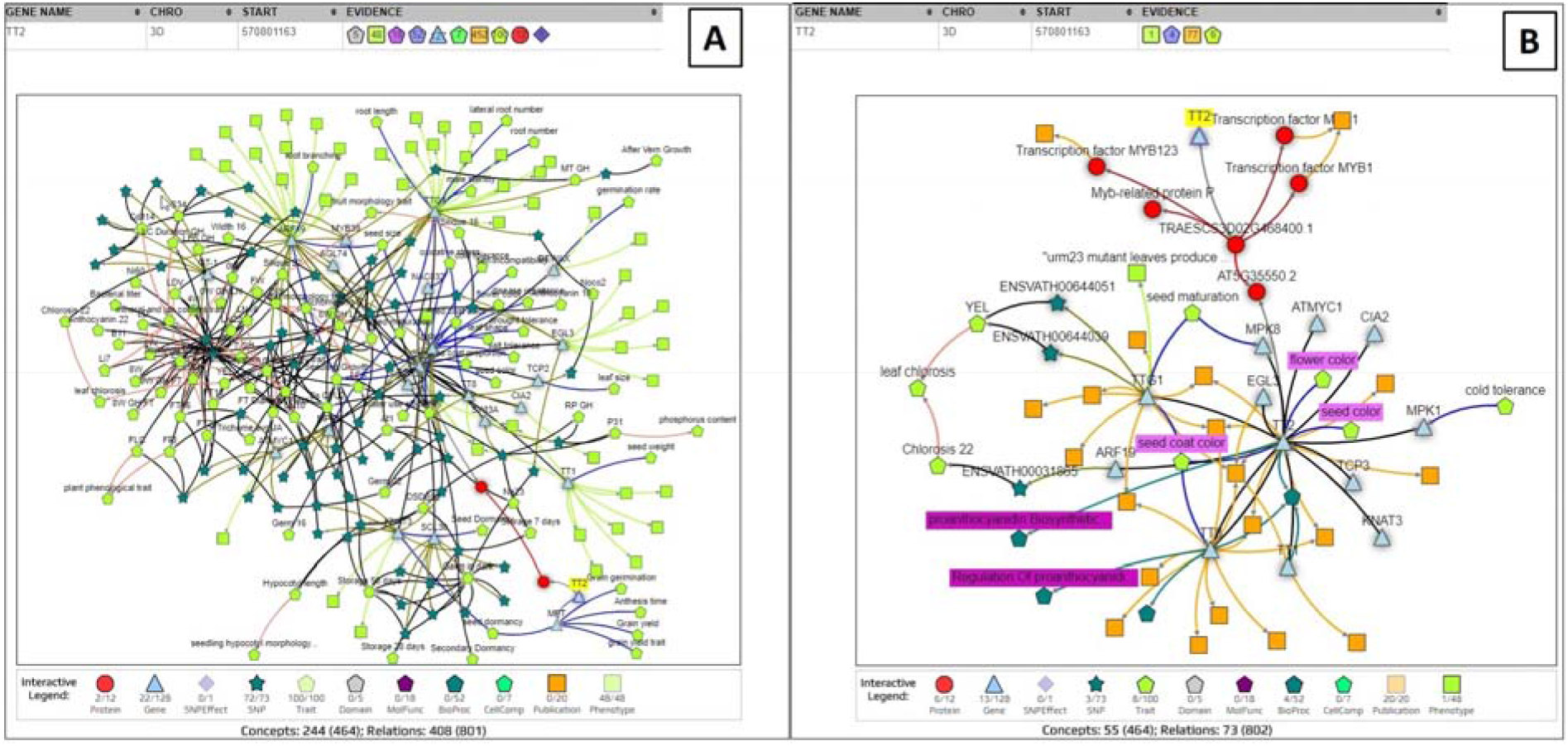
Gene View (top) and Network View (bottom) of KnetMiner. **(A)** Search results for TT2 only (without keywords). **(B)** Search results for TT2 and keywords for PHS and grain color.

To further refine and extend the search for evidence that links *TT2* to grain color and PHS, we can provide additional keywords relevant to the traits of interest. Seed germination and dormancy are the underlying developmental processes that activate or prevent pre-harvest sprouting in many grains and other seeds. The colour of the grain is known to be determined through accumulation of proanthocyanidin, an intermediate in the flavonoid pathway, found in the seed coat. These terms and phrases can be combined using boolean operators (AND, OR, NOT) and used in conjunction with a list of genes. Thus, we search for <u>TRAESCS3D02G468400</u> (*TT2*) and the keywords: *“seed germination” OR “seed dormancy” OR color OR flavonoid OR proanthocyanidin.* This time, KnetMiner filters the extracted *TT2* knowledge network (823 nodes) down to a smaller subgraph of 68 nodes and 87 relations in which every path from *TT2* to another node corresponds to a line of evidence to phenotype or molecular characteristics based on our keywords of interest (Figure 3B).

This auto-generated subgraph visualises complex information in a concise and connected format, helping facilitate biologically meaningful conclusions between *TT2* and phenotypes such PHS (see Supp Table 2). The subgraph indicates that *TT2* in wheat is predicted to regulate the transcriptional activation of *MFT*. It indicates that *MFT* has been linked in a recent publication to grain germination and seed dormancy in wheat (Nakamura S, n.d.; Zong Y, n.d.). It also reveals that the *MFT* ortholog in Arabidopsis is linked to decreased germination rate in the presence of ABA (Xi et al., 2010) and positive regulation of seed germination. To investigate potential links between grain color and other phenotypes, the TT2 network can be expanded with two clicks, to add interacting genes in wheat or model species along with their phenotypic information. For example, the Arabidopsis *TT2* ortholog is shown to interact with *TTG1* which has links to phenotypes such as lateral root number and root hair length in Arabidopsis (Bahmani R, n.d.; Bipei Zhang, 2017). Root hairs are tubular outgrowths from specific epidermal cells that function in nutrient and water absorption (Larry Peterson & Farquhar, 1996).

Overall the exploratory link analysis has generated a potential link between grain color and PHS due to *TT2-MFT* interaction and suggested a new hypothesis between two traits (PHS and root hair density) that were not part of the initial investigation and previously thought to be unrelated. Furthermore, it raises the possibility that *TT2* mutants might lead to increased root hairs and to higher nutrient and water absorption, and therefore cause early germination of the grain. More data and experiments will be needed to address this hypothesis and close the knowledge gap.

### Candidate gene prioritisation

Forward genetics studies, such as a genome-wide association study (GWAS) or quantitative trait loci (QTL) mapping, aim to identify regions in the genome where the genetic variation correlates with variation observed in a quantitative trait (e.g. general intelligence, days to flowering) (Atwell et al., 2010; Polderman et al., 2015; Sonah et al., 2015). They are based purely on statistical tests and do not take into account the biology in considering candidates. It is often difficult to elucidate which exact marker is biologically significant, particularly in the face of epistatic and epigenetic effects which are often not considered. GWAS and QTL regions can encompass many seemingly unrelated genes. Candidate gene analysis aims to identify the most likely cause for the phenotypic variation. The identification of candidate genes underlying QTL is not trivial, therefore genetic studies often stop after QTL mapping, or perform a basic search for genes with potentially interesting annotations.

For example, in a recent QTL study in Arabidopsis, a region on chromosome 4 was identified that contained overlapping QTLs for multiple petal traits (Abraham et al., 2013). As this QTL overlapped with the *ULTRAPETALA1* (*ULT1*) locus, a known floral meristem regulator with a role in petal development (Fletcher, 2001), the authors tested whether *ULT1* might be responsible for this QTL. However, the authors stated that among the ecotypes used in the study none showed any polymorphic sites within the *ULT1* coding or 2kb upstream region; and the T-DNA insertional mutation of *ULT1* showed no significant effect on petal form either. Taken together, the evidence suggested that *ULT1* was not responsible for the petal size QTL, and the causal gene remained unidentified as is the case in many other GWAS and QTL studies. Therefore, to explore this further, we analysed an overlapping petal size QTL (manuscript in preparation) using a more sophisticated and evidence-based search to see if the authors may have missed something. The biological processes underpinning the size of plant tissues and organs are likely to be related to changes on a cellular level. We therefore used as inputs to KnetMiner the location of a petal size QTL (chromosome 4, 9.92 - 10.18 Mb) and the keywords *“cell size” OR “cell cycle” OR “cell division”*. KnetMiner identified 71 genes in the QTL region and ranked them according to their relevance to the keywords (Figure 4A) (see Methods - Gene Ranking).

**Figure 4:**
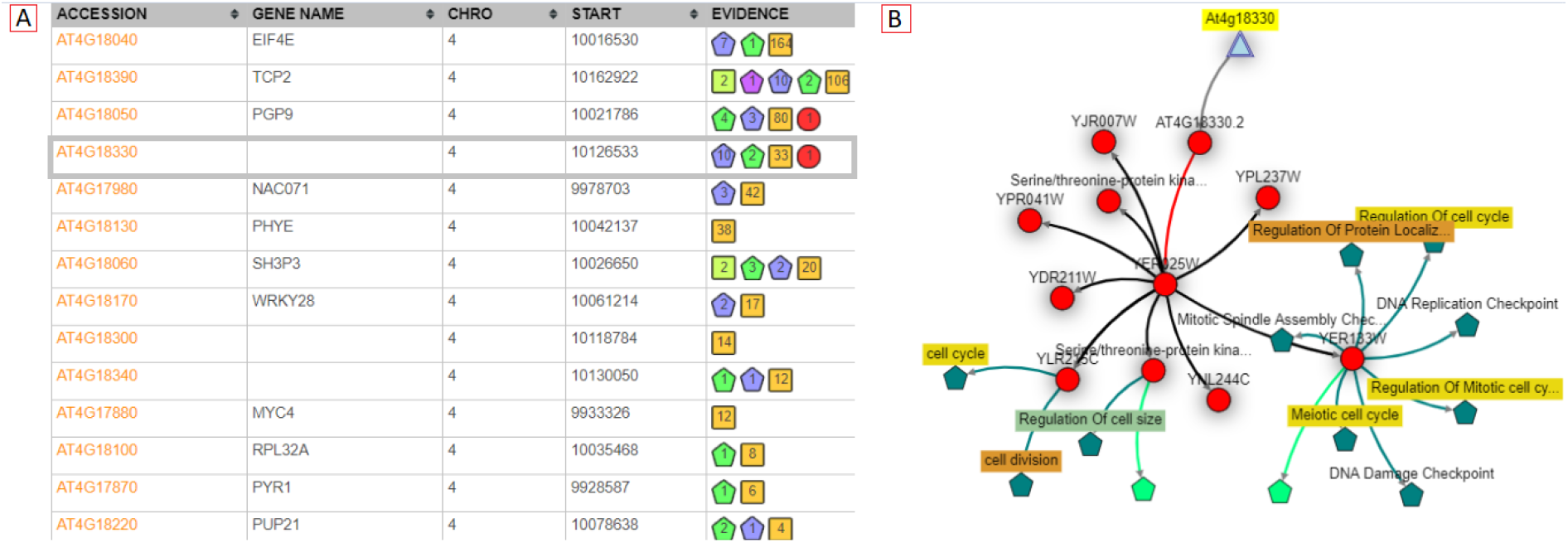
(**A**) Ranked list of genes shown in the Gene View. The Evidence column summarises the amount of related information within and across species. AT4G18330 is linked to 10 biological processes, 2 cellular components, 33 publications and 1 protein related to *“cell size” OR “cell cycle” OR “cell division”.* All linked publications are from the yeast ortholog. **(B)** Automatically generated subgraph for AT4G18330 and given keywords. The yeast ortholog YER025W interacts with several cell size, cell division and cell cycle related proteins.

The KnetMiner top 5 ranked genes included a poorly studied gene (AT4G18330) with no links to publications in Arabidopsis and a few high-level GO annotations. However, the KnetMiner subgraph for AT4G18330 indicated that the yeast ortholog YER025W (eIF-2-gamma) interacts with cell division cycle proteins such as CDC123 (Figure 4B). Although no knockouts were available for this gene, a polymorphism in the regulatory region was associated with altered cellular and petal phenotypes consistent with a role in petal size (manuscript in preparation). The ability to systematically and visually evaluate different layers of evidence arising from orthologs to interactions, is highly advantageous; it’s quick to view and as such, the most relevant genes can immediately be investigated further.

## METHODS

### Graph Pattern Mining

We have previously described our tools and methods to build FAIR genome-scale Knowledge Graphs (KG) using the KnetBuilder and rdf2neo data integration platforms (Brandizi et al., 2018a, 2018b; Hassani-Pak et al., 2016). Here we elaborate how KnetMiner uses the KG to extract biologically meaningful subgraphs that tell the story of complex traits and diseases. Biologically plausible patterns in the KG are collections of paths through the connected information that most biologists would generally agree to be informative when studying the function of a gene. Searching a KG for such patterns is akin to searching for relevant sentences containing evidence that supports a particular point of view within a book. Such evidence paths can be short e.g. Gene A was knocked out and phenotype *X* was observed; or alternatively the evidence path can be longer, e.g. Gene A in species *X* has an ortholog in species *Y*, which was shown to regulate the expression of a disease related gene (with a link to the paper). In the first example, the relationship between gene and disease is directly evident and experimentally proven, while in the second example the relationship is indirect and less certain but still biologically meaningful. There are many evidence types that should be considered for evaluating the relevance of a gene to a trait. In a KG context, a gene is considered to be, for example, related to ‘early flowering’ if any of its biologically plausible graph patterns contain nodes related to ‘early flowering’. In this context, the word ‘related’ doesn’t necessarily mean that the gene in question will have an effect on ‘flowering time’, but it means that there is a valid piece of evidence that a domain expert should consider when judging if the gene is related to ‘flowering time’.

We use the notion of a **semantic motif** to define a plausible path through the KG (Biemann et al., 2016). Our semantic motifs start with a gene node and end with other nodes representing biological entities, ontology terms, publications etc. For example, a path that travels from a Gene node to a GO-term, through an ortholog relation, is biologically plausible (orthologs have often the same function), while travelling through a paralog relation is not (paralogs often adapt new functions). KnetMiner instances can have a bespoke set of semantic motifs reflecting the data model of the KG built for a particular species or domain of interest. We are working towards migrating KnetMiner to support the Cypher graph query language and the Neo4j graph database as a practical and expressive way to define the graph searches that capture the semantic motifs of interest. Supp Table 3 contains example Cypher queries used in the public wheat KnetMiner along with summary statistics for each query. The KnetMiner gene search and subgraph generation are essentially based on these well-defined graph queries. Not every gene will necessarily match all semantic motifs, however, the ones it contains are extracted and their union is taken to produce a gene-centric subgraph (GCS). For example, the wheat KG has over 114,000 GCSs (one for each wheat gene) with sizes of min=1, max=6220 and mean=181 nodes.

Nodes that are included in a GCS are presumed to be transferable to the gene of interest, in contrast, concepts that are excluded from a GCS (although still part of the KG) are presumed to be irrelevant to the gene in question. Notably, if a semantic motif fails to capture an important biological motif, then downstream knowledge mining applications won’t be able to exploit this information.

### Graph Interestingness

Even a single GCS with hundreds of nodes can be complex and challenging to comprehend when shown to a user; let alone if combining GCSs for tens to hundreds of genes. There is therefore a need to filter and visualise the subset of information in the GCSs that is most interesting to a specific user. However, the interestingness of information is subjective and will depend on the biological question or the hypothesis that needs to be tested. A scientist with an interest in disease biology is likely to be interested in links to publications, pathways, and annotations related to diseases, while someone studying the biological process of grain filling is likely more interested in links to physiological or anatomical traits. To reduce information overload and visualise the most interesting pieces of information, we have devised two strategies. 1) In the case of a combined gene and keyword search, we use the keywords as a filter to show only paths in the GCS that connect genes with keyword related nodes, i.e. nodes that contain the given keywords in one of their node properties. In the special case where too many publications remain even after keyword filtering, we select the most recent N publications (default N=20). Nodes not matching the keyword are hidden but not removed from the GCS. 2) In the case of a simple gene query (without additional keywords), we initially show all paths between the gene and nodes of type phenotype/trait, i.e. any semantic motif that ends with a trait/phenotype, as this is considered the most important relationship to many KnetMiner users.

### Gene Ranking

We have developed a simple and fast algorithm to rank genes and their GCS for their importance. We give every node in the KG a weight composed of three components, referred to as SDR, standing for the **S**pecificity to the gene, **D**istance to the gene and **R**elevance to the search terms. **Specificity** reflects how specific a node is to a gene in question. For example, a publication that is cited (linked) by hundreds of genes receives a smaller weight than a publication which is linked to one or two genes only. We define the specificity of a node x as: 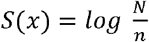 where n is the frequency of the node occurring in all N GCS. **Distance** assumes information which is associated more closely to a gene can generally be considered more certain, versus one that’s further away, e.g. inferred through homology and other interactions increases the uncertainty of annotation propagation. A short semantic motif is therefore given a stronger weight, whereas a long motif receives a weaker weight. Thus, we define the second weight as the inverse shortest path distance of a gene g and a node 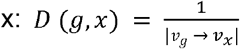. Both weights S and D are not influenced by the B X search terms and can therefore be pre-computed for every node in the KG. **Relevance** reflects the relevance or importance of a node to user-provided search terms using the well-established measure of inverse document frequency (IDF) and term frequency (TF) (Salton & Yang, 1973). TF*IDF forms the basis of the Lucene search engine library (https://lucene.apache.org/), used in KnetMiner. We define the relevance of node *x* to a search term *t* as *R*(*t,x*)= *TF x IDF* (*t,x*), where R=0 when no match is found and R=1 when the user does not provide any keywords. The three measures (S, D, and R) have unique and uncorrelated characteristics. Each node in KnetMiner is given a combined SDR weight. Therefore, for a given GCS *X*_*g*_ = {*x*, *x*_2_,…, *x*_*n*_} and search terms *t*, we define the *KnetScore* of a gene as:

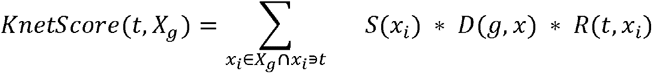

The sum considers only GCS nodes that contain the search terms. In the absence of search terms, we sum over all nodes of the GCS with R=1 for each node. The computation of the KnetScore (*SDR*-weights) requires graph traversals and string searches over the KG. Performing these operations on-the-fly would slow down the responsiveness of the application. Therefore at initialisation, KnetMiner pre-processes the KG and builds indices to speed up the *SDR* weight calculation. The pre-indexing time depends on a number of factors including number of available cores, the KG size, number of genes and number of semantic motifs. With the indices in place, the SDR-weight can be computed in constant time O(1). A KnetMiner search that returns n genes and m evidence nodes, can rank all genes in linear time O(n+m).

## DISCUSSION

Biological knowledge discovery is often hampered by the challenges of data integration and new approaches are needed to improve the efficiency, reproducibility, and objectivity of the process that leads to new ideas and hypotheses. KnetMiner provides a sophisticated search across a semantically rich knowledge graph built from large scale integration of public and private data sets. It addresses the needs of scientists who generally lack the time and the broad expertise that is necessary to connect, explore, and compare the wealth of genetic, ‘omics, and phenotypic information available in the literature and a wide range of related biological databases from key model and non-model species.

KnetMiner is commonly used by scientists in academia and industry to accelerate gene-trait discovery research. In several biological studies, KnetMiner enabled the identification of hidden relationships between important agronomic traits and potential candidate genes. The presented case studies have shown practical applications of KnetMiner to the understanding of challenging and complex traits in wheat and Arabidopsis. KnetMiner was used in 2014 to investigate traits such as height of biomass willows (Hanley & Karp, 2014) and has more recently become part of a wider roadmap for gene function characterization in crops (Adamski et al., 2020). Public KnetMiner resources (e.g. Arabidopsis, wheat, and rice) give a flavour of the capabilities that are in KnetMiner. While we have so far mostly concentrated on customising KnetMiner for plant sciences and crop improvement, the software we have developed is generic and KGs and KnetMiner can readily be built for other species. Compared to biological discovery platforms available for specific species (Carvalho-Silva et al., 2019; Miller et al., 2017; Mungall et al., 2017), KnetMiner is species-agnostic and therefore provides a more cost-effective delivery platform for application to new species. KnetMiner is available as a Docker image from DockerHub and can easily be deployed with a provided sample KG.

Different KnetMiner views for exploring the search output have been developed; each view has a different aim and helps address different questions. The main design principle was to divide the visualisation into two steps. First, to present the results in formats that are intuitive and familiar to biologists, such as tables and chromosome views, allowing them to explore the data, make choices as to which gene to view, or refine the query if needed. These initial views help users to reach a certain level of confidence with the selection of potential candidate genes. However, they do not tell the biological story that links candidate genes to traits and diseases. In a second step, to enable the stories and their evidence to be investigated in full detail, the Network View visualises highly complex information in a concise and connected format, helping facilitate biologically meaningful conclusions. Consistent graphical symbols are used for representing evidence types throughout the different views, so that users develop a certain level of familiarity, before being exposed to networks with complex interactions and rich content.

The methods (graph pattern mining, graph interestingness and gene ranking) that power the KnetMiner user interface are also available as API calls and can be used to embed visualisations of gene-centric subgraphs in third party web applications or to integrate graph analytics and gene ranking in custom workflows. For example, the KnetMiner REST API is used in Ensembl Plants (Bolser et al., 2017), The Triticeae Toolbox (Blake et al., 2016) and GrainGenes (Blake et al., 2019) to link gene sequences to rich gene knowledge graphs. The graph database backend, as well as the FAIR-based data management policies, are another development in which we are investing our efforts, which have the main advantage of allowing us to build a data asset that has the potential to be useful to a wealth of applications, complementary to KnetMiner. The SPARQL and Cypher endpoints have the benefit of providing a layer of access to data that have a more general use than gene-centric knowledge exploration and which, for instance, could be obtained with scripts accessing APIs, workflow tools like Galaxy (Afgan et al., 2018), or data analytics workbenches like Jupyter (Kluyver et al., 2016). This is facilitated by adhering to the well-known good practice of the FAIR principles, which includes the adoption of common data schemas and ontologies (Garcia et al., 2017).

## CONCLUSION

Scientists spend a considerable amount of time searching for new clues and ideas by synthesizing many different sources of information and using their expertise to generate hypotheses. KnetMiner is a user-friendly platform for biological knowledge discovery and exploratory data mining. It allows humans and machines to effectively connect the dots in life science data and literature, search the connected data in an innovative way, and then return the results in an accessible, explorable, yet concise format that can be easily interrogated to generate new insights. We have developed KnetMiner knowledge graphs and applications for a range of species including plants, crops, insects, pathogens, livestock and even a Human SARS-CoV-2 knowledge graph to help investigate Covid-19. We are beginning to explore new use cases of KnetMiner to crop improvement and breeding, microbial ecology, pathogen-host interaction and other domains. We are rapidly improving the usability of the software, adding new features and extending the knowledge mining approaches. The latest version of the KnetMiner software and documentation is available at: https://knetminer.org

## DECLARATIONS

### Availability of data and materials

Project name: KnetMiner - Knowledge Network Miner

Project home page: https://knetminer.org

Source code: https://github.com/Rothamsted/knetminer

Docker image: https://hub.docker.com/r/knetminer/knetminer

Deployment instructions: https://github.com/Rothamsted/knetminer/wiki/

Knowledge Graph Endpoints: http://knetminer.org/data

Operating system(s): Platform independent

Programming language: Java and JavaScript

Other requirements: Docker

License: MIT

Any restrictions to use by non-academics: database licence needed

### Competing interests

The authors declare that they have no competing interests.

### Funding

This work was supported by the UKRI Biotechnology and Biological Sciences Research Council (BBSRC) through the Designing Future Wheat ISP (BB/P016855/1), DiseaseNetMiner TRDF (BB/N022874/1), ONDEX SABR funding (BB/F006039/1) and National Capability in Crop Phenotyping (BB/J004464/1). CR, KHP, AS are additionally supported by strategic funding to Rothamsted Research from BBSRC. JHD also acknowledges support from the National Science Foundation (cROP project 1340112).

### Authors’ contributions

KHP designed the approach as part of his dissertation with CR, collected results, and drafted the manuscript. KHP, AS, MB, JH and the KnetMiner team implemented the KnetMiner framework and maintain its public instances. SA helped to build the Arabidopsis and wheat knowledge graphs. AP and JHD provided the biological use cases. All authors read, reviewed and approved the final manuscript.

## Acknowledgements

We acknowledge all the past and present members of the KnetMiner Bioinformatics team at Rothamsted for their scientific inputs, software testing and technical support: Emma Bailey, Dan Smith, Robert King, David Hughes, Monika Mistry, Minja Zorc, Fengyuan Hu, Jan Taubert, William Brown and Ricardo Gregorio. We acknowledge all our collaborators who contributed to the development of the KnetMiner resources and software in the past including Martin Castellote, Maria Esch, Vasiliki Koutra, Haolin Li, Philipp Bayer, Ramil Mauleon, Cristobal Uauy, Jean-Luc Jannink, Clay Birkett, Uwe Schulz, Steve Hanley, Francis Newson and Richard Holland.

## Notes

https://knetminer.org

https://knetminer.org/Triticum_aestivum/

https://knetminer.org/Arabidopsis_thaliana/

https://knetminer.org/data

https://hub.docker.com/r/knetminer/knetminer

https://github.com/Rothamsted/knetminer/blob/master/species/wheat-beta/ws/cypher-queries.txt

### Competing Interest Statement

The authors have declared no competing interest.

### Summary of Updates

Updated the title and abstract to better communicate the main advances of KnetMiner. Updated the Data and Materials availability statement.

## REFERENCES

Abraham, M. C., Metheetrairut, C., & Irish, V. F. (2013). Natural variation identifies multiple loci controlling petal shape and size in Arabidopsis thaliana. PloS One, 8(2), e56743.

Adamski, N. M., Borrill, P., Brinton, J., Harrington, S. A., Marchal, C., Bentley, A. R., Bovill, W. D., Cattivelli, L., Cockram, J., Contreras-Moreira, B., Ford, B., Ghosh, S., Harwood, W., Hassani-Pak, K., Hayta, S., Hickey, L. T., Kanyuka, K., King, J., Maccaferrri, M., … Uauy, C. (2020). A roadmap for gene functional characterisation in crops with large genomes: Lessons from polyploid wheat. eLife, 9. https://doi.org/10.7554/eLife.55646

Afgan, E., Baker, D., Batut, B., van den Beek, M., Bouvier, D., Cech, M., Chilton, J., Clements, D., Coraor, N., Grüning, B. A., Guerler, A., Hillman-Jackson, J., Hiltemann, S., Jalili, V., Rasche, H., Soranzo, N., Goecks, J., Taylor, J., Nekrutenko, A., & Blankenberg, D. (2018). The Galaxy platform for accessible, reproducible and collaborative biomedical analyses: 2018 update. Nucleic Acids Research, 46(W1), W537–W544.

Alabdullah, A. K., Borrill, P., Martin, A. C., Ramirez-Gonzalez, R. H., Hassani-Pak, K., Uauy, C., Shaw, P., & Moore, G. (2019). A Co-Expression Network in Hexaploid Wheat Reveals Mostly Balanced Expression and Lack of Significant Gene Loss of Homeologous Meiotic Genes Upon Polyploidization. Frontiers in Plant Science, 10, 1325.

Ali, M., Hoyt, C. T., Domingo-Fernández, D., Lehmann, J., & Jabeen, H. (n.d.). BioKEEN: A library for learning and evaluating biological knowledge graph embeddings. https://doi.org/10.1101/475202

Alshahrani, M., & Hoehndorf, R. (2018). Semantic Disease Gene Embeddings (SmuDGE): phenotype-based disease gene prioritization without phenotypes. Bioinformatics, 34(17), i901–i907.

Atwell, S., Huang, Y. S., Vilhjálmsson, B. J., Willems, G., Horton, M., Li, Y., Meng, D., Platt, A., Tarone, A. M., Hu, T. T., Jiang, R., Wayan Muliyati, N., Zhang, X., Amer, M. A., Baxter, I., Brachi, B., Chory, J., Dean, C., Debieu, M., … Nordborg, M. (2010). Genome-wide association study of 107 phenotypes in *Arabidopsis thaliana* inbred lines. Nature, 465(7298), 627.

Bahmani R, E. al. (n.d.). The Density and Length of Root Hairs Are Enhanced in Response to Cadmium and Arsenic by Modulating Gene Expressions Involved in Fate Determination … - PubMed - NCBI. Retrieved September 3, 2018, from https://www.ncbi.nlm.nih.gov/pubmed/27933081

Biemann, C., Chris, B., Lachezar, K., Stefanie, R., & Karsten, W. (2016). Network Motifs Are a Powerful Tool for Semantic Distinction. In Understanding Complex Systems (pp. 83–105).

Bipei Zhang, A. S. (2017). TRANSPARENT TESTA GLABRA 1-Dependent Regulation of Flavonoid Biosynthesis. Plants, 6(4). https://doi.org/10.3390/plants6040065

Blake, V. C., Birkett, C., Matthews, D. E., Hane, D. L., Bradbury, P., & Jannink, J.-L. (2016). The Triticeae Toolbox: Combining Phenotype and Genotype Data to Advance Small-Grains Breeding. The Plant Genome, 9(2). https://doi.org/10.3835/plantgenome2014.12.0099

Blake, V. C., Woodhouse, M. R., Lazo, G. R., Odell, S. G., Wight, C. P., Tinker, N. A., Wang, Y., Gu, Y. Q., Birkett, C. L., Jannink, J.-L., Matthews, D. E., Hane, D. L., Michel, S. L., Yao, E., & Sen, T. Z. (2019). GrainGenes: centralized small grain resources and digital platform for geneticists and breeders. Database: The Journal of Biological Databases and Curation, 2019. https://doi.org/10.1093/database/baz065

Bolser, D. M., Staines, D. M., Perry, E., & Kersey, P. J. (2017). Ensembl Plants: Integrating Tools for Visualizing, Mining, and Analyzing Plant Genomic Data. Methods in Molecular Biology, 1533, 1–31.

Boyle, E. A., Li, Y. I., & Pritchard, J. K. (2017). An Expanded View of Complex Traits: From Polygenic to Omnigenic. Cell, 169(7), 1177–1186.

Brandizi, M., Singh, A., Rawlings, C., & Hassani-Pak, K. (2018a). Towards FAIRer Biological Knowledge Networks Using a Hybrid Linked Data and Graph Database Approach. Journal of Integrative Bioinformatics. https://doi.org/10.1515/jib-2018-0023

Brandizi, M., Singh, A., Rawlings, C., & Hassani-Pak, K. (2018b). Getting the best of Linked Data and Property Graphs: rdf2neo and the KnetMiner Use Case. SWAT4LS Proceedings. https://doi.org/10.6084/m9.figshare.7314323.v1

Carvalho-Silva, D., Pierleoni, A., Pignatelli, M., Ong, C., Fumis, L., Karamanis, N., Carmona, M., Faulconbridge, A., Hercules, A., McAuley, E., Miranda, A., Peat, G., Spitzer, M., Barrett, J., Hulcoop, D. G., Papa, E., Koscielny, G., & Dunham, I. (2019). Open Targets Platform: new developments and updates two years on. Nucleic Acids Research, 47(D1), D1056–D1065.

De Bie, T. (2013). Subjective Interestingness in Exploratory Data Mining. In Lecture Notes in Computer Science (pp. 19–31).

De Bie, T., & Spyropoulou, E. (2013). A Theoretical Framework for Exploratory Data Mining: Recent Insights and Challenges Ahead. In Lecture Notes in Computer Science (pp. 612–616).

Ehrlinger, L., & Wöß, W. (2016). Towards a Definition of Knowledge Graphs. SEMANTiCS (Posters, Demos, SuCCESS), 48. https://www.researchgate.net/profile/Wolfram_Woess/publication/323316736_Towards_a_Definition_of_Knowledge_Graphs/links/5a8d6e8f0f7e9b27c5b4b1c3/Towards-a-Definition-of-Knowledge-Graphs.pdf

Fensel, D., Şimşek, U., Angele, K., Huaman, E., Kärle, E., Panasiuk, O., Toma, I., Umbrich, J., & Wahler, A. (2020). Introduction: What Is a Knowledge Graph? In Knowledge Graphs (pp. 1–10). Springer, Cham.

Fletcher, J. C. (2001). The ULTRAPETALA gene controls shoot and floral meristem size in Arabidopsis. Development, 128(8), 1323–1333.

Fox, P., & Hendler, J. (2011). Changing the equation on scientific data visualization. Science, 331(6018), 705–708.

Garcia, L., Giraldo, O., Garcia, A., & Dumontier, M. (2017). Bioschemas: schema. org for the Life Sciences. Proceedings of SWAT4LS. http://ceur-ws.org/Vol-2042/paper33.pdf

Hanley, S. J., & Karp, A. (2014). Genetic strategies for dissecting complex traits in biomass willows (Salix spp.). Tree Physiology, 34(11), 1167–1180.

Harrington, S. A., Backhaus, A. E., Singh, A., & Hassani-Pak, K. (2019). Validation and characterisation of a wheat GENIE3 network using an independent RNA-Seq dataset. bioRxiv. https://www.biorxiv.org/content/10.1101/684183v1.abstract

Hassani-Pak, K., Castellote, M., Esch, M., Hindle, M., Lysenko, A., Taubert, J., & Rawlings, C. (2016). Developing integrated crop knowledge networks to advance candidate gene discovery. Applied & Translational Genomics, 11, 18–26.

Hassani-Pak, K., Legaie, R., Canevet, C., van den Berg, H. A., Moore, J. D., & Rawlings, C. J. (2010). Enhancing data integration with text analysis to find proteins implicated in plant stress response. Journal of Integrative Bioinformatics, 7(3). https://doi.org/10.2390/biecoll-jib-2010-121

Holmes, J. H. (2014). Knowledge Discovery in Biomedical Data: Theory and Methods. In Methods in Biomedical Informatics (pp. 179–240).

Holzinger, A., & Jurisica, I. (2014). Knowledge Discovery and Data Mining in Biomedical Informatics: The Future Is in Integrative, Interactive Machine Learning Solutions. In Lecture Notes in Computer Science (pp. 1–18).

Isenberg, T., Isenberg, P., Chen, J., Sedlmair, M., & Möller, T. (2013). A Systematic Review on the Practice of Evaluating Visualization. IEEE Transactions on Visualization and Computer Graphics, 19(12), 2818–2827.

Kluyver, T., Ragan-Kelley, B., Pérez, F., Granger, B. E., Bussonnier, M., Frederic, J., Kelley, K., Hamrick, J. B., Grout, J., Corlay, S., Ivanov, P., Avila, D., Abdalla, S., & Willing, C. (2016). Jupyter Notebooks - a publishing format for reproducible computational workflows. https://doi.org/10.3233/978-1-61499-649-1-87

Larry Peterson, R., & Farquhar, M. L. (1996). Root hairs: Specialized tubular cells extending root surfaces. The Botanical Review; Interpreting Botanical Progress, 62(1), 1–40.

Lee, B., Isenberg, P., Riche, N. H., & Carpendale, S. (2012). Beyond Mouse and Keyboard: Expanding Design Considerations for Information Visualization Interactions. IEEE Transactions on Visualization and Computer Graphics, 18(12), 2689–2698.

Messina, A., Fiannaca, A., La Paglia, L., La Rosa, M., & Urso, A. (2018). BioGraph: a web application and a graph database for querying and analyzing bioinformatics resources. BMC Systems Biology, 12(Suppl 5), 98.

Miller, J., Town, C., Stuerzlinger, W., & Provart, N. J. (2017). ePlant: visualizing and exploring multiple levels of data for hypothesis generation in plant biology. The Plant. http://www.plantcell.org/content/29/8/1806.short

Mohamed, S. K., Nováček, V., & Nounu, A. (2019). Discovering Protein Drug Targets Using Knowledge Graph Embeddings. Bioinformatics. https://doi.org/10.1093/bioinformatics/btz600

Mungall, C. J., McMurry, J. A., Köhler, S., Balhoff, J. P., Borromeo, C., Brush, M., Carbon, S., Conlin, T., Dunn, N., Engelstad, M., Foster, E., Gourdine, J. P., Jacobsen, J. O. B., Keith, D., Laraway, B., Lewis, S. E., NguyenXuan, J., Shefchek, K., Vasilevsky, N., … Haendel, M. A. (2017). The Monarch Initiative: an integrative data and analytic platform connecting phenotypes to genotypes across species. Nucleic Acids Research, 45(D1), D712–D722.

Nakamura S, E. al. (n.d.). A wheat homolog of MOTHER OF FT AND TFL1 acts in the regulation of germination. - PubMed - NCBI. Retrieved August 30, 2018, from https://www.ncbi.nlm.nih.gov/pubmed/21896881

Nilsson-Ehle, H. (1914). Zur Kenntnis der mit der keimungsphysiologie des weizens in zusammenhang stehenden inneren faktoren.

Pavelin, K., Cham, J. A., de Matos, P., Brooksbank, C., Cameron, G., & Steinbeck, C. (2012). Bioinformatics meets user-centred design: a perspective. PLoS Computational Biology, 8(7), e1002554.

Polderman, T. J. C., Benyamin, B., de Leeuw, C. A., Sullivan, P. F., van Bochoven, A., Visscher, P. M., & Posthuma, D. (2015). Meta-analysis of the heritability of human traits based on fifty years of twin studies. Nature Genetics, 47(7), 702–709.

Russell-Rose, T., Chamberlain, J., & Azzopardi, L. (2018). Information retrieval in the workplace: A comparison of professional search practices. Information Processing & Management, 54(6), 1042–1057.

Sacchi, L., & Holmes, J. H. (2016). Progress in Biomedical Knowledge Discovery: A 25-year Retrospective. Yearbook of Medical Informatics, Suppl 1, S117–S129.

Salton, G., & Yang, C. S. (1973). On the Specification of Term Values in Automatic Indexing. Journal of Documentation.

Sears, E. R. (1944). Cytogenetic Studies with Polyploid Species of Wheat. II. Additional Chromosomal Aberrations in Triticum Vulgare. Genetics, 29(3), 232.

Sheth, A., Padhee, S., & Gyrard, A. (2019). Knowledge Graphs and Knowledge Networks: The Story in Brief. IEEE Internet Computing, 23(4), 67–75.

Singh, A., Rawlings, C. J., & Hassani-Pak, K. (2018). KnetMaps: a BioJS component to visualize biological knowledge networks. F1000Research, 7, 1651.

Sonah, H., O’Donoughue, L., Cober, E., Rajcan, I., & Belzile, F. (2015). Identification of loci governing eight agronomic traits using a GBS-GWAS approach and validation by QTL mapping in soya bean. In Plant Biotechnology Journal (Vol. 13, Issue 2, pp. 211–221). https://doi.org/10.1111/pbi.12249

Stephens, Z. D., Lee, S. Y., Faghri, F., Campbell, R. H., Zhai, C., Efron, M. J., Iyer, R., Schatz, M. C., Sinha, S., & Robinson, G. E. (2015). Big Data: Astronomical or Genomical? PLoS Biology, 13(7), e1002195.

Sweis, B. M., Abram, S. V., Schmidt, B. J., Seeland, K. D., MacDonald, A. W., 3rd, Thomas, M. J., & Redish, A. D. (2018). Sensitivity to “sunk costs” in mice, rats, and humans. Science, 361(6398), 178–181.

The International Wheat Genome Sequencing Consortium (IWGSC), IWGSC RefSeq principal investigators:, Appels, R., Eversole, K., Feuillet, C., Keller, B., Rogers, J., Stein, N., IWGSC whole-genome assembly principal investigators:, Pozniak, C. J., Choulet, F., Distelfeld, A., Poland, J., Ronen, G., Sharpe, A. G., Whole-genome sequencing and assembly:, Pozniak, C., Barad, O., Baruch, K., … Manuscript writing team: (2018). Shifting the limits in wheat research and breeding using a fully annotated reference genome. Science, 361(6403), eaar7191.

Xiaoxue, L., Xuesong, B., Longhe, W., Bingyuan, R., Shuhan, L., & Lin, L. (2019). Review and Trend Analysis of Knowledge Graphs for Crop Pest and Diseases. IEEE Access, 7, 62251–62264.

Xi, W., Liu, C., Hou, X., & Yu, H. (2010). MOTHER OF FT AND TFL1 regulates seed germination through a negative feedback loop modulating ABA signaling in Arabidopsis. The Plant Cell, 22(6), 1733–1748.

Yoon, B.-H., Kim, S.-K., & Kim, S.-Y. (2017). Use of Graph Database for the Integration of Heterogeneous Biological Data. Genomics & Informatics, 15(1), 19–27.

Zong Y, E. al. (n.d.). Allelic Variation and Transcriptional Isoforms of Wheat TaMYC1 Gene Regulating Anthocyanin Synthesis in Pericarp. - PubMed - NCBI. Retrieved August 30, 2018, from https://www.ncbi.nlm.nih.gov/pubmed/28983311

