## Supplemental Table 1 for "KnetMiner: a comprehensive approach for supporting evidence-based gene discovery and complex trait analysis across species"

**Supplementary Table 1:** Information types and instances in the wheat *TT2 (*TRAESCS3D02G468400) gene-centric subgraph.

| **Information type** | **Instances** |
|---|---:|
| Publication | 452 |
| Gene | 128 |
| Trait | 101 |
| SNP | 73 |
| Biological Process (GO) | 52 |
| Phenotype | 48 |
| Molecular Function (GO) | 18 |
| Protein | 12 |
| Cellular Component (GO) | 7 |
| Protein Domain | 5 |
| SNP Effect | 1 |
