## Supplemental Table 2 for "KnetMiner: a comprehensive approach for supporting evidence-based gene discovery and complex trait analysis across species"

**Supplementary Table 2:** Examples of relation types and properties in the keyword-filtered TT2 *(*TRAESCS3D02G468400) subgraph

| **Node A** | **Node B** | **Relation Type** | **Relation Properties** |
|---|---|---|---|
| TT2-3D | MFT-3B | regulates | p-value=0.01; evidence:GENIE3; data=850_samples_wheat_rna_seq |
| TT2-3D | TT2-3B | homoeolog | Ensembl Compara |
| MFT-3B | Grain germination (CO:0000011) | co-occurs | Recent studies in both Arabidopsis and wheat have uncovered a new role of MOTHER OF FT AND TFL1 (MFT) in seed germination. (PMID: 24932489) |
| MFT-3B | Seed dormancy (TO:0000253) | co-occurs | Mapping analysis showed that MFT on chromosome 3A (MFT-3A) colocalized with the seed dormancy quantitative trait locus (QTL) QPhs.ocs-3A (PMID :21896881) |
| AtMFT | Decreased germination rate | has_phenotype | Decreased rate of germination in the presence of ABA. (PMID:20551347) |
| AtMFT | Positive regulation of seed germination (GO:0010030) | participates_in | Inferred from Biological aspect of Ancestor<br>Inferred from Mutant Phenotype |
| AtTT2 | AtTTG1 | interacts | Two-hybrid (PMID: 15255866) |
| AtTTG1 | Lateral root number (TO:0001013) | co-occurs | Moreover, transgenic TTG1-overexpression (TTG1-OX) seedlings exhibited enhanced root length and lateral root number compared to wild-type seedlings grown under normal or stress conditions. (PMID: 23306631) |
| TT2 | TT8 (bHLH) | interacts | Recently the TaMYC1 gene encoding bHLH transcription factor has been isolated from the bread wheat (Triticum aestivum L.) genome and shown to co-locate with the Pp3 gene conferring purple pericarp color. (PMID: 28983311) |
