## Supplemental Table 3 for "KnetMiner: a comprehensive approach for supporting evidence-based gene discovery and complex trait analysis across species"

**Supplementary Table 3:** Examle of semantic motfics (in Cypher language) used in KnetMiner with number of matches found in the Wheat Knowledge Graph (Release 45)

| Query | Paths in KG | Paths per Gene | Path Length |
|---|---|---|---|
| `(Gene)-[:enc]->(Protein)-[:ortho]-(Protein)<-[:enc]-(Gene)-[:genetic\|physical]-(Gene)-[:pub_in]->(Publication)` | 9,549,126 | 87.6 | 11 |
| `(Gene)-[:homoeolog\|regulates\|genetic\|physical]-(Gene)-[:has_mutant\|has_variation]->(SNP)-[:leads_to]->(SNPEffect)` | 3,478,148 | 32.4 | 7 |
| `(Gene)-[:part_of]->(CoExpCluster)-[:enriched_for]->(PlantOntologyTerm)` | 2,734,304 | 25.1 | 5 |
| `(Gene)-[:homoeolog\|regulates\|genetic\|physical]-(Gene)` | 2,273,339 | 21.0 | 3 |
| `(Gene)-[:homoeolog\|regulates\|genetic\|physical]-(Gene)-[:occ_in]->(Publication)` | 2,040,168 | 18.7 | 5 |
| `(Gene)-[:part_of]->(CoExpCluster)-[:enriched_for]->(BioProc)` | 1,751,039 | 16.2 | 5 |
| `(Gene)-[:enc]->(Protein)-[:ortho]-(Protein)<-[:enc]-(Gene)-[:genetic\|physical]-(Gene)-[:participates_in]->(BioProc)` | 1,497,522 | 13.7 | 11 |
| `(Gene)-[:enc]->(Protein)-[:ortho]-(Protein)<-[:enc]-(Gene)-[:genetic\|physical]-(Gene)-[:has_variation]->(SNP)-[:associated_with]->(Trait)-[:pub_in]->(Publication)` | 1,329,197 | 12.2 | 15 |
| `(Gene)-[:enc]->(Protein)-[:ortho]-(Protein)<-[:enc]-(Gene)-[:genetic\|physical]-(Gene)-[:located_in]->(CelComp)` | 1,133,500 | 10.5 | 11 |
| `(Gene)-[:enc]->(Protein)-[:ortho]-(Protein)<-[:enc]-(Gene)-[:genetic\|physical]-(Gene)-[:has_variation]->(SNP)-[:associated_with]->(Trait)-[:is_part_of]-(Trait)` | 1,062,833 | 9.8 | 15 |
| `(Gene)-[:enc]->(Protein)-[:ortho]-(Protein)<-[:enc]-(Gene)-[:genetic\|physical]-(Gene)-[:has_function]->(MolFunc)` | 1,051,504 | 9.7 | 11 |

<table>
<tr><td>(Gene)-[:enc]->(Protein)-[:ortho]-(Protein)<-[:enc]-(Gene)-[:pub_in]->(Publication)</td><td>994,535</td><td>9.1</td><td>9</td></tr>
<tr><td>(Gene)-[:enc]->(Protein)-[:h_s_s|ortho|xref*0..1]-(Protein)-[:pub_in]->(Publication)</td><td>859,379</td><td>7.9</td><td>7</td></tr>
<tr><td>(Gene)-[:enc]->(Protein)-[:h_s_s|ortho|xref*0..1]-(Protein)-[:has_function]->(MolFunc)</td><td>738,183</td><td>6.8</td><td>7</td></tr>
<tr><td>(Gene)-[:enc]->(Protein)-[:h_s_s|ortho|xref*0..1]-(Protein)-[[:participates_in]->(BioProc)</td><td>589,120</td><td>5.4</td><td>7</td></tr>
<tr><td>(Gene)-[:enc]->(Protein)-[:h_s_s|ortho|xref*0..1]-(Protein)-[:has_domain]->(ProtDomain)</td><td>581,162</td><td>5.3</td><td>7</td></tr>
<tr><td>(Gene)-[:enc]->(Protein)-[:h_s_s|ortho|xref*0..1]-(Protein)</td><td>557,435</td><td>5.1</td><td>5</td></tr>
<tr><td>(Gene)-[:enc]->(Protein)-[:h_s_s|ortho|xref*0..1]-(Protein)-[:located_in]->(CelComp)</td><td>488,173</td><td>4.5</td><td>7</td></tr>
<tr><td>(Gene)-[:enc]->(Protein)-[:ortho]-(Protein)<-[:enc]-(Gene)-[:genetic|physical]-(Gene)-[:has_observ_pheno]->(Phenotype)</td><td>484,240</td><td>4.4</td><td>11</td></tr>
<tr><td>(Gene)-[:enc]->(Protein)-[:ortho]-(Protein)<-[:enc]-(Gene)-[:genetic|physical]-(Gene)-[:has_variation]->(SNP)-[[:associated_with]->(Trait)</td><td>481,474</td><td>9.3</td><td>13</td></tr>
<tr><td>(Gene)-[:homoeolog|regulates|genetic|physical]-(Gene)-[:part_of]->(Path)</td><td>450,971</td><td>4.1</td><td>5</td></tr>
<tr><td>(Gene)-[:enc]->(Protein)-[:ortho]-(Protein)<-[:enc]-(Gene)-[:genetic|physical]-(Gene)-[:cooc_wi]-(Trait)</td><td>429,313</td><td>4.0</td><td>11</td></tr>
<tr><td>(Gene)-[:enc]->(Protein)-[:ortho]-(Protein)<-[:enc]-(Gene)-[:genetic|physical]-(Gene)</td><td>390,646</td><td>3.6</td><td>9</td></tr>
<tr><td>(Gene)-[:enc]->(Protein)-[:ortho]-(Protein)<-[:enc]-(Gene)-[:has_variation]->(SNP)-[[:associated_with]->(Trait)</td><td>334,404</td><td>3.1</td><td>11</td></tr>
<tr><td>(Gene)-[:enc]->(Protein)-[:h_s_s|ortho|xref*0..1]-(Protein)-[:cat_c]->(EC)</td><td>235,234</td><td>2.2</td><td>7</td></tr>
</table>

<table>
<tr><td>(Gene)-[:enc]->(Protein)-[:ortho]-(Protein)<-[:enc]-(Gene)-[:has_variation]->(SNP)-[:associated_with]->(Trait)-[:pub_in]->(Publication)</td><td>224,482</td><td>2.1</td><td>13</td></tr>
<tr><td>(Gene)-[:homoeolog|regulates|genetic|physical]-(Gene)-[:cooc_wi]-(Trait)</td><td>223,087</td><td>2.1</td><td>5</td></tr>
<tr><td>(Gene)-[:homoeolog|regulates|genetic|physical]-(Gene)-[:inv_in]->(Reaction)</td><td>222,101</td><td>2.0</td><td>5</td></tr>
<tr><td>(Gene)-[:enc]->(Protein)-[:ortho]-(Protein)<-[:enc]-(Gene)-[:located_in]->(CelComp)</td><td>201,102</td><td>1.8</td><td>9</td></tr>
<tr><td>(Gene)-[:enc]->(Protein)-[:ortho]-(Protein)<-[:enc]-(Gene)-[:participates_in]->(BioProc)</td><td>199,505</td><td>1.8</td><td>9</td></tr>
<tr><td>(Gene)-[:enc]->(Protein)-[:ortho]-(Protein)<-[:enc]-(Gene)-[:has_variation]->(SNP)-[:associated_with]->(Trait)-[:is_part_of]-(Trait)</td><td>176,308</td><td>1.6</td><td>13</td></tr>
<tr><td>(Gene)-[:enc]->(Protein)-[:ortho]-(Protein)<-[:enc]-(Gene)-[:has_function]->(MolFunc)</td><td>174,136</td><td>1.6</td><td>9</td></tr>
<tr><td>(Gene)-[:has_mutant|has_variation]->(SNP)-[:leads_to]-> (SNPEffect)</td><td>137,057</td><td>1.3</td><td>5</td></tr>
<tr><td>(Gene)-[:enc]->(Protein)</td><td>133,346</td><td>1.2</td><td>3</td></tr>
<tr><td>(Gene)-[:part_of]->(CoExpCluster)-[:part_of]->(CoExpStudy)</td><td>103,429</td><td>1.0</td><td>5</td></tr>
<tr><td>(Gene)-[:enc]->(Protein)-[:h_s_s|ortho|xref*0..1]-(Protein)-[:cat_c]->(EC)-[:equ]-(MolFunc)</td><td>102,225</td><td>0.9</td><td>9</td></tr>
<tr><td>(Gene)-[:occ_in]->(Publication)</td><td>94,752</td><td>0.9</td><td>3</td></tr>
<tr><td>(Gene)-[:enc]->(Protein)-[:ortho]-(Protein)<-[enc_10_8_d:enc]-(gene_8:Gene)</td><td>70,891</td><td>0.7</td><td>7</td></tr>
<tr><td>(Gene)-[:enc]->(Protein)-[:h_s_s|ortho|xref*0..1]-(Protein)-[:is_a]->(Enzyme)</td><td>54,192</td><td>0.5</td><td>7</td></tr>
<tr><td>(Gene)-[:homoeolog|regulates|genetic|physical]-(Gene)-[:has_mutant|has_variation]->(SNP)-[:associated_with]-> (Trait)</td><td>53,805</td><td>0.5</td><td>7</td></tr>
<tr><td>(Gene)-[:enc]->(Protein)-[:h_s_s|ortho|xref*0..1]-(Protein)-[:xref]-(Protein)</td><td>48,366</td><td>0.4</td><td>7</td></tr>
</table>

| | | | |
|---|---|---|---|
| (Gene)-[:enc]->(Protein)-[:ortho]-(Protein)<-[:enc]-(Gene)-[:has_observ_pheno]->(Phenotype) | 45,438 | 0.4 | 9 |
| (Gene)-[:enc]->(Protein)-[:ortho]-(Protein)<-[:enc]-(Gene)-[:cooc_wi]-(Trait) | 40,389 | 0.4 | 9 |
| (Gene)-[:part_of]->(Path) | 14,213 | 0.1 | 3 |
| (Gene)-[:inv_in]->(Reaction) | 6,274 | 0.1 | 3 |
| (Gene)-[:cooc_wi]-(Trait) | 5,976 | 0.1 | 3 |
| (Gene)-[:has_mutant\|has_variation]->(SNP)-[:associated_with]->(Trait) | 1,501 | 0.0 | 5 |
